# Genome sequence of a diabetes-prone desert rodent reveals a mutation hotspot around the ParaHox gene cluster

**DOI:** 10.1101/093401

**Authors:** Adam D Hargreaves, Long Zhou, Josef Christensen, Ferdinand Marlétaz, Shiping Liu, Fang Li, Peter Gildsig Jansen, Enrico Spiga, Matilde Thye Hansen, Signe Vendelbo Horn Pedersen, Shameek Biswas, Kyle Serikawa, Brian A Fox, William R Taylor, John F Mulley, Guojie Zhang, R Scott Heller, Peter W H Holland

**Author notes:** Joint corresponding authors.

## Abstract

The sand rat *Psammomys obesus* is a gerbil native to deserts of North Africa and the Middle East^1^. Sand rats survive with low caloric intake and when given high carbohydrate diets can become obese and develop type II diabetes^2^ which, in extreme cases, leads to pancreatic failure and death^3,4^. Previous studies have reported inability to detect the *Pdx1* gene or protein in gerbils^5–7^, suggesting that absence of this key insulin-regulating homeobox gene might underlie diabetes susceptibility. Here we report sequencing of the sand rat genome and discovery of an extensive, mutationally-biased GC-rich genomic domain encompassing many essential genes, including the elusive *Pdx1.* The sequence of *Pdx1* has been grossly affected by GC-biased mutation leading to the highest divergence observed in the animal kingdom. In addition to molecular insights into restricted caloric intake in a desert species, the discovery that specific chromosomal regions can be subject to elevated mutation rate has widespread significance to evolution.

---

Linking molecular change to phenotypic change is a central goal of evolutionary biology. Adaptation to arid environments is particularly interesting because of the extreme physiological demands imposed by low food and water availability. The sand rat *Psammomys obesus* (Fig. 1a) is a member of the subfamily Gerbillinae, most species of which live in deserts and arid environments (Fig. 1b). *P. obesus* has emerged as a model for research into diet-induced type II diabetes because, if provided with high carbohydrate diets, the majority of individuals become obese and develop classic diabetes symptoms, in the most extreme cases leading to pancreatic failure and death^2,3^.

**Figure 1.**
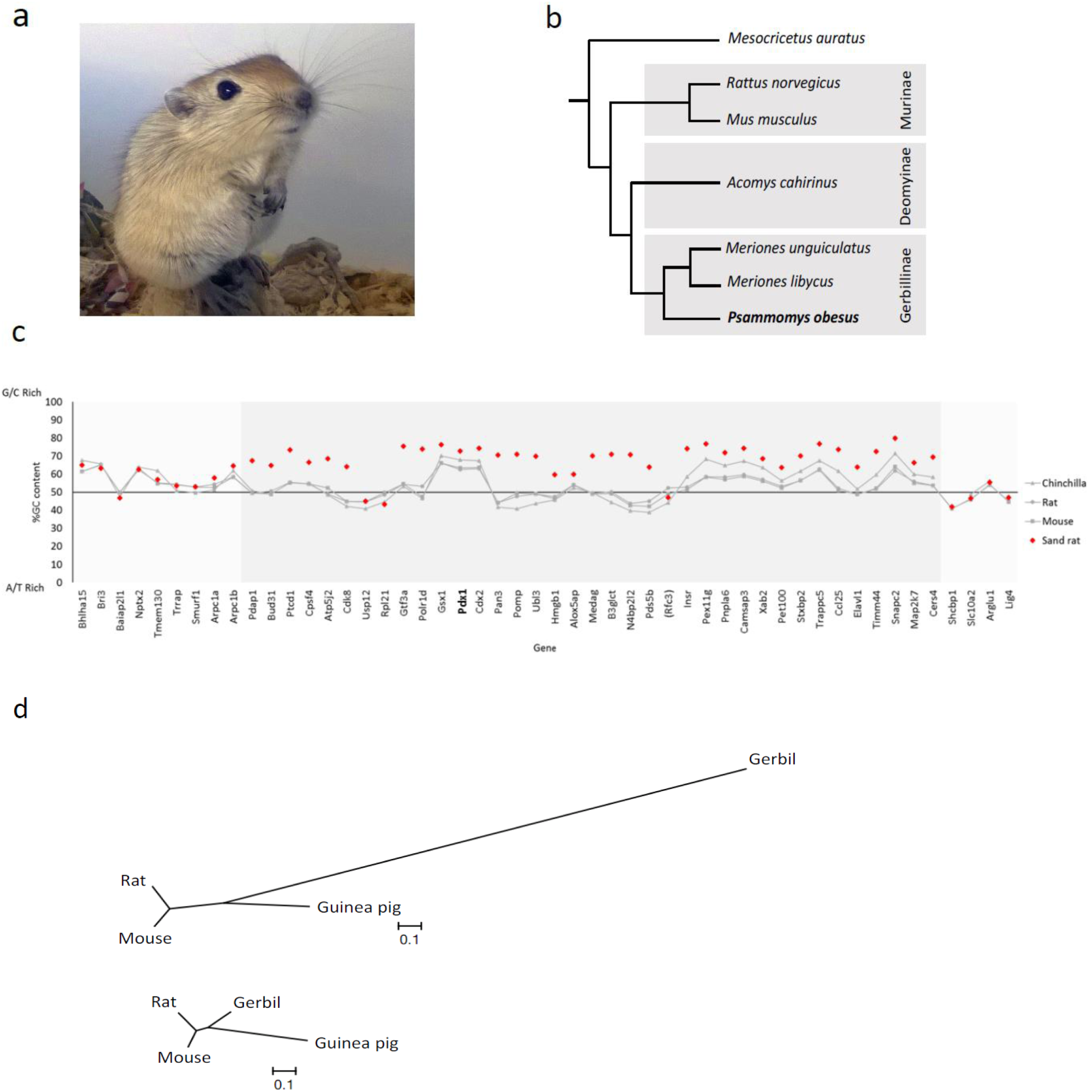
The sand rat and its genomic hotspot of mutation. (a) Juvenile sand rat *Psammomys obesus.* (b) Cladogram of representative murid rodents indicating the phylogenetic position of sand rat. (c) GC content of genes around the ParaHox cluster of sand rat and other rodents *(Mus musculus, Rattus norvegicus, Chinchilla lanigera)* revealing a chromosomal hotspot of GC skew in sand rat (shaded in grey). Genes shown in inferred ancestral gene order; parentheses around *Rfc3* indicate this gene has been transposed to a different genomic location in sand rat. Sand rat GC values based on transcriptome and genome sequences; when partial only alignable sequence is compared. (d) Unrooted phylogenetic trees inferred from synonymous changes (dS) only from concatenated alignments of 26 genes in the mutational hotspot (top) and 100 random genes (bottom).

In searching for the molecular basis of this unusual phenotype, attention has been paid to the *Pdx1* homeobox gene, also called *Ipf1, Idx1, Stf1* or *Xlox*^8–12^, the central and most highly conserved member of the ParaHox gene cluster^13^. *Pdx1* is the only member of the Pdx gene family in tetrapods, and encodes a homeodomain that has been invariant across their evolution. Mammalian *Pdx1* is expressed in pancreatic beta-cells^14^ and encodes a homeodomain transcription factor that acts as a transcriptional activator of *insulin* and other pancreatic hormone genes^15,16^. A pivotal role in insulin regulation is also reflected in the association of heterozygous *Pdx1* mutations with maturity-onset diabetes of the young *(MODY4)* and type II diabetes mellitus in humans^16^. Contrary to the usual conservation, several studies have reported inability to detect *Pdx1* in gerbils, including *P. obesus,* by immunocytochemistry, Western blotting or PCR^5–7^, leading to the hypothesis that the gene has been lost, compromising ability to regulate insulin. Such a conclusion would raise further questions, since in addition to its adult functions, *Pdx1* is also essential for pancreatic development in the embryo. For example, targeted deletion in mice causes loss of pancreas and anterior duodenum and is lethal^8,17^. In humans, pancreatic agenesis has been reported in a patient with a homozygous frameshift mutation before the *Pdx1* homeobox, and in a compound heterozygous patient with substitution mutations in helices 1 and 2 of the homeodomain^18–20^.

To resolve the conundrum of a putatively absent ‘essential’ gene, we sequenced the *P. obesus* genome using a standard shotgun strategy (Illumina), using a combination of short and long insert libraries, initially at 85.5X coverage (Supplementary information section 1). This assembly lacked a *Pdx1* gene supporting the prevailing hypothesis of a loss of the *Pdx1* gene in gerbils. However, a synteny comparison between *P. obesus* and other mammals delineated a contiguous block of 88 genes (Supplementary table S2.3.1) missing from the assembly including several genes essential to basic cellular functions, such as *Brca2* and *Cdk8,* in addition to *Pdx1.* This led us to suspect that standard short read sequencing may have given an incomplete genome assembly. To resolve whether this represented a large-scale deletion or an unusual genomic region, we sequenced the transcriptomes of *P. obesus* liver, pancreatic islets and duodenum, which strikingly contained transcripts for many of the missed genes (Supplementary information section 2; Supplementary table S2.3.1). Furthermore, these transcripts show unusually high GC content in most cases, indicating that a large contiguous stretch of elevated GC had either been under-represented in initial sequencing data or had failed to assemble correctly, most likely due to nucleotide compositional bias. We term such cryptic or hidden sequence ‘dark DNA’. We therefore isolated GC-rich *P. obesus* genomic DNA by Caesium Chloride gradient centrifugation, sequenced this fraction after limited amplification using Illumina MiSeq overlapping paired-end reads, and re-assembled the genome incorporating this sequence data (Supplementary information section 1.5). This gave a refined assembly with a total size of 2.38 Gb and a scaffold N50 of 10.4 Mb (Table 1; Supplementary information sections 1,3,4,6), including much of the ‘dark DNA’ region in several scaffolds, and contains genes syntenic to a region of chromosome 12 in rat and a region of chromosome 5 and the subtelomeric region of chromosome 8 in mouse. Comparison of GC content between species demonstrates that sand rat genes are elevated in GC content across this chromosomal region, syntenic to 12 Mb of the rat genome (Fig. 1c; Supplementary information section 9). This large region encompasses a 250 kb repeat-rich scaffold containing the sand rat ParaHox cluster and its well-characterised genomic neighbours. We inferred a high W (weak, A/T) to S (strong, G/C) allelic mutation rate in this region of the *P. obesus* genome when compared with randomly selected genomic regions or homologous regions in other species of rodent (Fig 1d; Supplementary information section 11, Supplementary tables S11.1 & S11.2). The existence of a localised GC-biased stretch of the *P. obesus* genome is striking and of far-reaching importance, and implies the existence of elevated and biased mutational pressure, acting in one region of the genome.

**Table 1.**
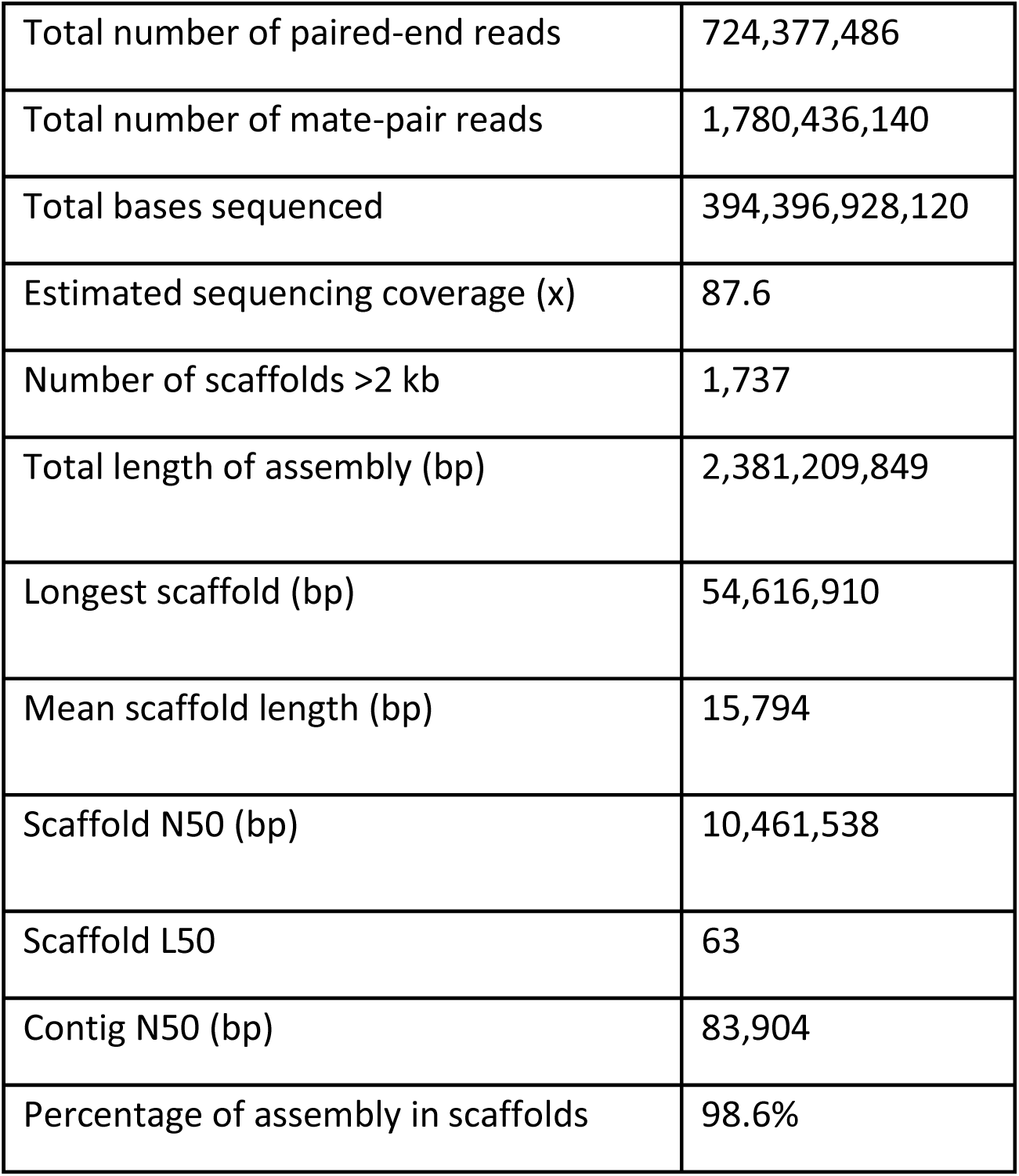
Metrics of sand rat raw genomic sequencing data and final genome assembly. Coverage was calculated using an estimated genome size of 2.51 Gb based on a k-mer analysis (Supplementary information section 1.3) and is based upon paired-end sequencing data only.

|  |  |
| --- | --- |
| Total number of paired-end reads | 724,377,486 |
| Total number of mate-pair reads | 1,780,436,140 |
| Total bases sequenced | 394,396,928,120 |
| Estimated sequencing coverage (x) | 87.6 |
| Number of scaffolds >2 kb | 1,737 |
| Total length of assembly (bp) | 2,381,209,849 |
| Longest scaffold (bp) | 54,616,910 |
| Mean scaffold length (bp) | 15,794 |
| Scaffold N50 (bp) | 10,461,538 |
| Scaffold L50 | 63 |
| Contig N50 (bp) | 83,904 |
| Percentage of assembly in scaffolds | 98.6% |

The full coding sequence of the *P. obsesus Pdx1* gene was deduced from the refined genome and transcriptome assemblies, and the gene was found to be expressed in sand rat pancreatic islets and duodenum (Supplementary information section 7). The 60 amino acid homeodomain of Pdx1 shows 100% conservation across other mammals for which data are available; however, in *P. obesus* there are a remarkable 15 amino acid differences in the homeodomain, making this by far the most divergent *Pdx1* gene discovered in the animal kingdom (Fig. 2a). All but one of the amino acid changes are caused by A/T to G/C mutation. The N-terminal and C-terminal regions are also divergent with numerous deletions, although the hexapeptide motif used in heterodimer formation with TALE proteins is conserved (Fig. 2b). Despite its radical divergence, *Pdx1* is the closest homeodomain by blastp and phylogenetic analysis places it as a rodent *Pdx1* on a long branch (Fig. 2c); extensive synteny with the ParaHox region of mouse and rat confirms it is the true and single *Pdx1* ortholog (Supplementary table S7.1). Evidence that the locus is functional includes expression in pancreas and duodenum, and the fact that extensive polymorphism is found in the 3‘ untranslated region but is very limited in the coding sequence (Supplementary information section 11), indicating that the coding region is under functional constraint despite extensive mutation. Extreme deviation from the expected sequence explains why antibodies and PCR failed to detect *Pdx1* in sand rat^5–7^.

**Figure 2.**
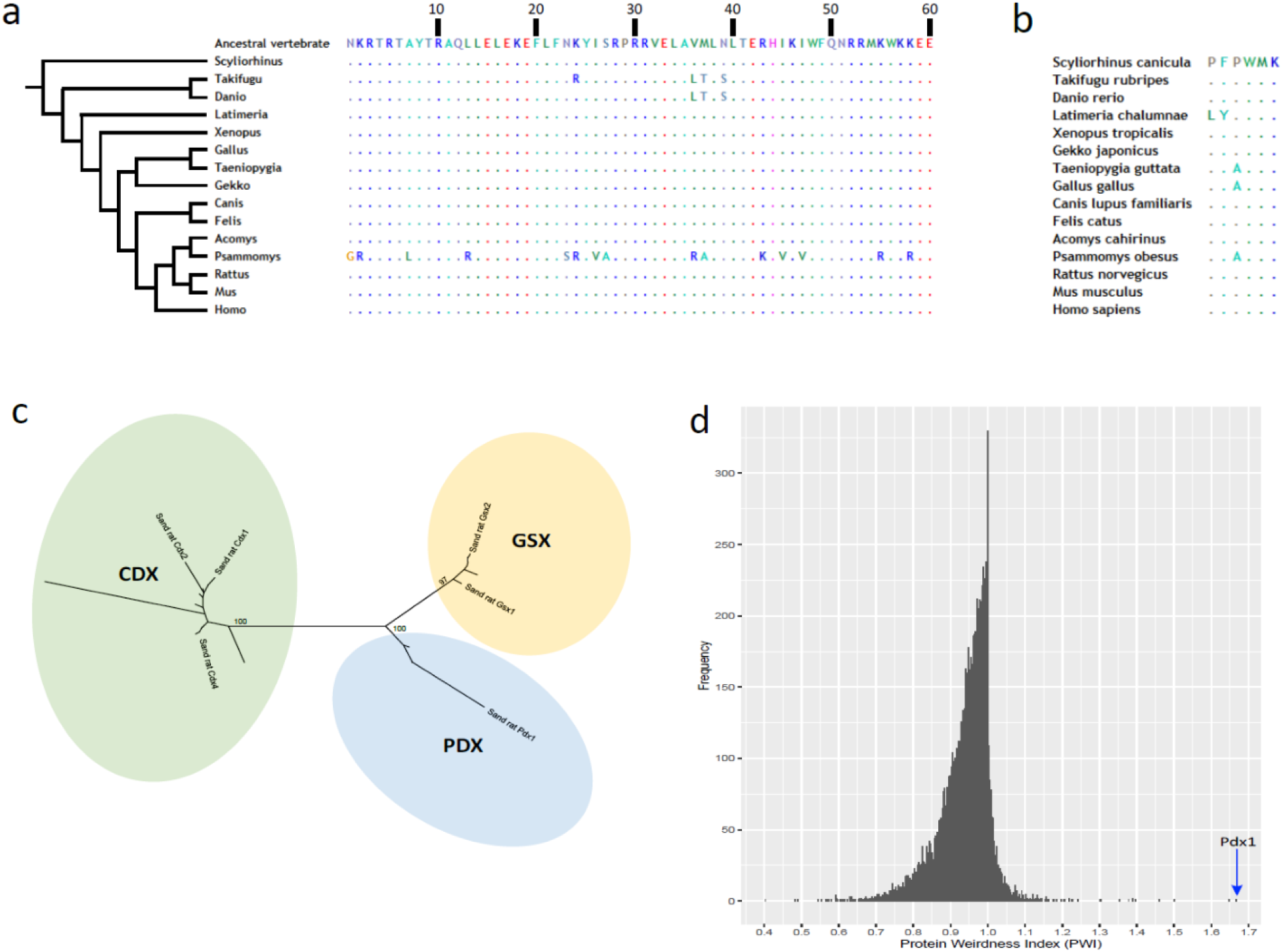
Molecular divergence of sand rat Pdx1. (a) Alignment of Pdx1 homeodomain sequences across vertebrates. (b) Alignment of Pdx1 hexapeptide domain across vertebrates. (c) Maximum likelihood tree of ParaHox proteins showing divergent *Psammomys obesus* Pdx1; species included are sand rat, mouse, zebra finch, spotted gar, amphioxus (full tree Figure S7.1). (d) Histogram of Protein Weirdness Index (PWI) values for 1:1:1 mammalian orthologs of the sand rat predicted proteins: Pdx1 is marked by an arrow.

These findings indicate that GC-biased mutation has driven radical changes in an otherwise highly conserved homeobox gene; these changes could be maladaptive and constrain the physiological capability of the sand rat, or adaptive enhancing ability to live in arid regions. To test if the extent of sequence divergence is unusual for sand rat proteins, we calculated a ‘protein weirdness index’ (PWI) (Supplementary information section 5) for all 1:1 mammalian orthologs by dividing mouse-human protein sequence identity by mouse-sand rat sequence identity (Fig. 2d). This is distinct from identifying the fastest evolving proteins, and specifically identifies proteins that have undergone uncharacteristic divergence in sand rat. We find the majority of sand rat proteins are highly similar to mouse or human (mode PWI = 1.0); in contrast, Pdx1 is unusually divergent (mouse-sand rat 54.82%, mouse-human 91.37%; PWI = 1.67). To test if other genes implicated in glucose metabolism or pancreatic function are also divergent, we compiled a list of 45 candidates from human studies including all genes implicated in monogenic diabetes^21^ and genes for which coding sequence variants have been strongly associated with T2D^22^. Of the 33 genes with clear 1:1:1 orthologs between human, mouse and sand rat, 32 lie between position 225 and 10,195 in our PWI ranking, indicating that they are not unusually divergent in sand rat. Pdx1 is ranked 1st and is the most unusually divergent protein identified in the sand rat predicted proteome (Supplementary information section 8; Supplementary table S5.1).

The mutations fixed in sand rat *Pdx1* gene do not cause frameshifts or truncations in known domains, and molecular modelling reveals that the sand rat Pdx1 homeodomain has the ability to form all three helices required for DNA binding (Fig. 3a). To examine if these mutations have resulted in subtle effects on the stability of DNA binding we deployed molecular dynamics simulations with atomistic representation of Pdx1 homeodomains, DNA target and solvent. From the post-processing of the molecular dynamics simulations we estimated the enthalpy of binding between sand rat and mouse (or other mammal) Pdx1 and monomer DNA binding sites using the MM-PBSA (Molecular Mechanics Poisson Boltzmann Surface Area) method (Supplementary information section 10). Target DNA sequences used were core Pdx1-binding sites of the mouse *insulin* A1 promoter and its sand rat ortholog. From 200 ns molecular dynamics simulations the enthalpy of binding for protein-DNA interaction was calculated to be lower for sand rat than for mouse Pdx1 (mean −140 kcal/mol vs. mean −122 kcal/mol), indicative of sand rat Pdx1 binding DNA more ‘tightly’ than is normal for the mammalian Pdx1 protein (Fig. 3b). One amino acid change was responsible for much of the difference: a Leu to Arg substitution in alpha helix 1 (homeodomain position 13), leading to the positive side chain of Arg making a new indirect contact with the phosphate backbone of DNA. A second substitution, Val to Arg in alpha helix 2 (homeodomain position 36), makes a smaller contribution (Fig. 3c). We also detect modifications to specific base interactions, with sand rat residues Met54 and Arg58 making new contacts to A and T bases within the TAAT core. Hence, stronger DNA binding is most likely driven by increased contacts with the backbone of DNA, coupled with decreased sequence-specificity of DNA interaction. These results suggest that sand rat Pdx1 is suboptimal in DNA-binding affinity and specificity.

**Figure3.**
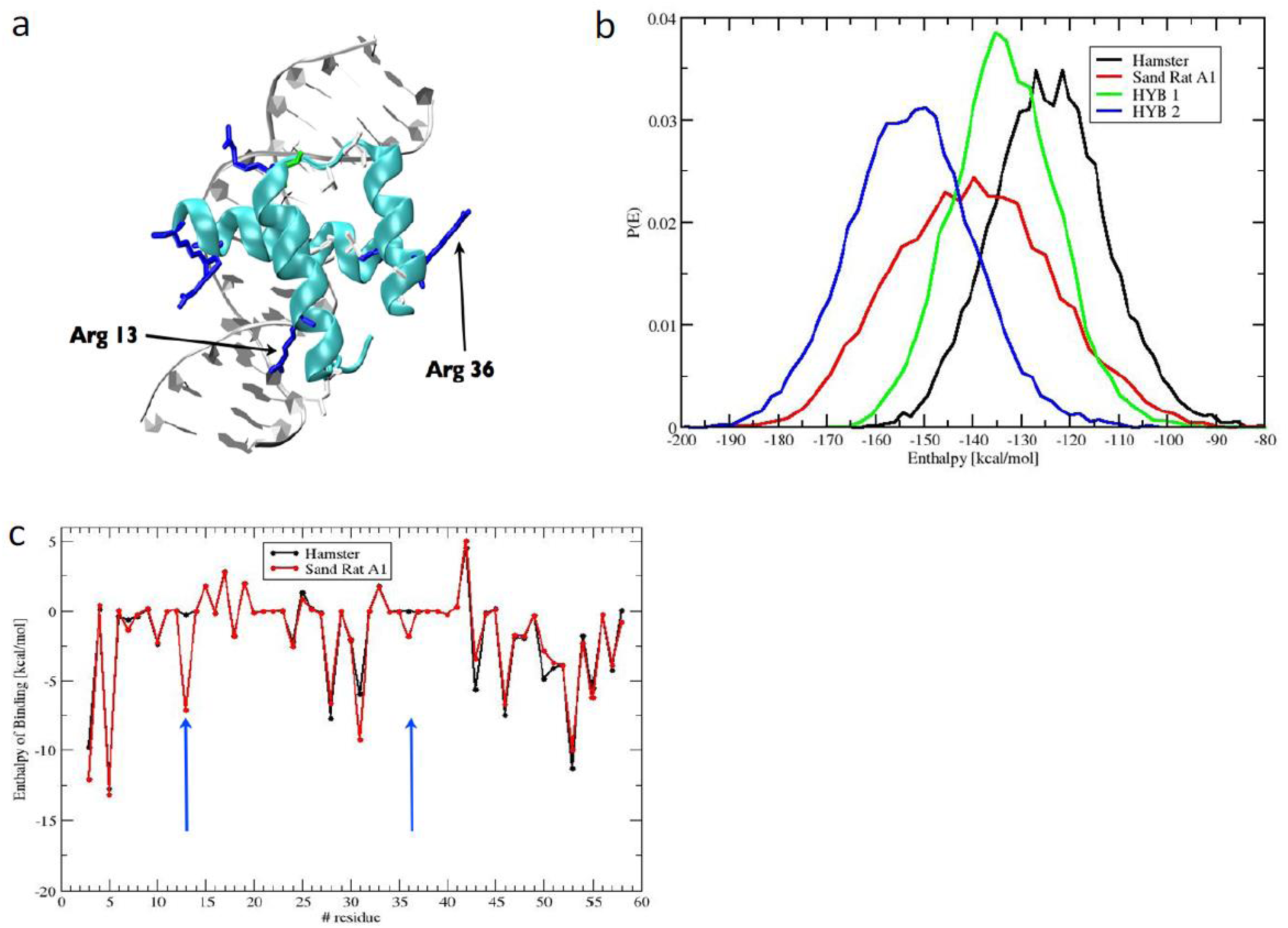
Molecular modelling of sand rat Pdx1 binding. (a) Molecular model of sand rat Pdx1 homeodomain bound to DNA. The two amino acid changes indicated are the largest contributors to altered enthalpy of binding. (b) Probability distributions of the enthalpy of binding of homeodomain protein-DNA interactions between hamster (normal vertebrate) Pdx1/hamster insulin A1 DNA element (black), sand rat Pdx1/sand rat A1 element (red), hamster Pdx1/sand rat A1 (green) and sand rat Pdx1/hamster A1 (blue) inferred by molecular dynamics simulations and MM-PBSA; sand rat Pdx1 homeodomain has the lowest enthalpy of binding (higher affinity) for each DNA target. (c) Per-site enthalpy of binding comparison between hamster and sand rat Pdx1 revealing contribution of amino acid changes at homeodomain positions 13 and 36 to reduced enthalpy of binding (higher affinity).

We conclude that an unusual genomic region of biased mutation arose in the evolutionary lineage of the sand rat. One consequence of this hotspot of mutation was the generation of GC-bias in the *Pdx1* gene of *P. obesus;* this forced modification of the Pdx1 protein sequence, affecting its ability to regulate insulin gene transcription and most likely transcription of other pancreatic genes. The sand rat Pdx1 hexapeptide, which mediates co-factor interactions^23^, is intact, which may explain why pancreatic development proceeds permitting viable sand rat embryogenesis. We suggest mutation-driven changes have played a role in constraining or adapting the sand rat, and possibly other gerbil species, to arid environments and low caloric intake. Biased gene conversion is a known mechanism that causes GC-biased mutation^24,25^; hence we suggest this mechanism, driven by elevated localised recombination, is generating a hotspot of skewed base composition. The genomic region we describe here was not detected by standard sequencing approaches, raising the possibility that other such dark DNA regions could be widespread features of animal genomes, thus far largely overlooked in comparative animal genomics. Indeed, GC-rich genes are also missing from the chicken genome assembly^26,27^. Hotspots of mutation could drive rapid evolutionary change at the molecular level, and it will be important to decipher to what extent such hotspots have constrained and influenced evolutionary adaptation across the animal kingdom.

## Author contributions

ADH and PWHH conducted GC-rich DNA isolation, sequencing and analysis; LZ, SL, FL and GZ performed genome assembly and annotation; MTH prepared DNA samples; SVHP and ADH extracted RNA samples; KS, SB, BF and ADH performed RNA-seq and assembly; JC performed laboratory investigations underpinning subsequent work; ADH, LZ, PGJ, JFM and FM undertook bioinformatic analyses; ES and WRT ran molecular dynamic simulations; RSH, GZ and PWHH initiated and directed the research; PWHH, ADH, LZ, GZ and RSH drafted the manuscript. All authors approved the final manuscript.

## Acknowledgements

This work was funded principally by the European Research Council under the European Union's Seventh Framework Programme (FP7/2007-2013 ERC grant 268513 awarded to PWHH), a Strategic Priority Research Program of the Chinese Academy of Sciences (XDB13000000 awarded to GZ) and Novo Nordisk A/S (coordinated by RSH). ES and WRT were supported by the Francis Crick Institute under awards: FC001179. The Crick receives its core funding from Cancer Research UK,the UK Medical Research Council, and the Wellcome Trust. We thank Natasha Ng, Gemma Marfany, Thomas Dunwell, Fei Xu, Shan Quah, Anna Gloyn, Christine Hirschberger, Juliane Cohen, Rhys Morgan, Lorna Witty, Monica Martinez Alonso and Thomas Brekke for assistance and advice, and the Oxford Genomics Centre for GC-rich sequencing.

